# Hyperedge bundling: A practical solution to spurious interactions in MEG/EEG source connectivity analyses

**DOI:** 10.1101/219311

**Authors:** Sheng H. Wang, Muriel Lobier, Felix Siebenhühner, Tuomas Puoliväli, Satu Palva, J. Matias Palva

**Author notes:** The correspondence should be addressed to: Sheng H., Wang, Neuroscience Center, University of Helsinki or J. Matias Palva, Neuroscience Center, University of Helsinki.

## Abstract

Inter-areal functional connectivity (FC), neuronal synchronization in particular, is thought to constitute a key systems-level mechanism for coordination of neuronal processing and communication between brain regions. Evidence to support this hypothesis has been gained largely using invasive electrophysiological approaches. In humans, neuronal activity can be non-invasively recorded only with magneto- and electroencephalography (MEG/EEG), which have been used to assess FC networks with high temporal resolution and whole-scalp coverage. However, even in source-reconstructed MEG/EEG data, signal mixing, or “source leakage”, is a significant confounder for FC analyses and network localization.

Signal mixing leads to two distinct kinds of false-positive observations: artificial interactions (AI) caused directly by mixing and spurious interactions (SI) arising indirectly from the spread of signals from true interacting sources to nearby false loci. To date, several interaction metrics have been developed to solve the AI problem, but the SI problem has remained largely intractable in MEG/EEG all-to-all source connectivity studies. Here, we advance a novel approach for correcting SIs in FC analyses using source-reconstructed MEG/EEG data.

Our approach is to bundle observed FC connections into hyperedges by their adjacency in signal mixing. Using realistic simulations, we show here that bundling yields hyperedges with good separability of true positives and little loss in the true positive rate. Hyperedge bundling thus significantly decreases graph noise by minimizing the false-positive to true-positive ratio. Finally, we demonstrate the advantage of edge bundling in the visualization of large-scale cortical networks with real MEG data. We propose that hypergraphs yielded by bundling represent well the set of true cortical interactions that are detectable and dissociable in MEG/EEG connectivity analysis.

**Highlights:**

- A true interaction often is “ghosted” into a multitude of spurious edges (SI)
- Effective in controlling and illustrating SI
- Hyperedges have much improved TPR and graph quality
- Advantages in visualizing connectivity

## 1 Introduction

Large-scale neuronal networks, *e.g.*, manifested by functional, directed, and effective connectivity(Karl J. 2011), are thought to be critical for healthy brain functions while their abnormalities are thought to underlie many brain diseases (Brookes et al., 2016; Bullmore and Sporns 2009; Bullmore and Sporns 2012; Fornito et al., 2015; Papo et al., 2014; Petersen and Sporns 2015; Rubinov 2015; Sporns 2014; Uhlhaas and Singer 2010; Uhlhaas and Singer 2006). Currently, magneto-and electro-encephalography (MEG/EEG) are the only noninvasive electrophysiological tools for studying connectivity networks with millisecond-range temporal resolution and good coverage of the cortical surface (Kujala et al., 2008; Palva and Palva 2012; S. Baillet et al., 2001; Salmelin and Baillet 2009). Accurately identifying interaction dynamics from MEG/EEG data is of crucial importance for understanding their role in human cognition and its deficits.

To date, numerous interaction metrics have been developed and utilized to assess functional connectivity (FC) in terms of amplitude-, phase-, and phase-amplitude correlations within or across frequency bands for pairs of electrophysiological signals (Bastos and Schoffelen 2016; Kreuz 2011; O’Neill et al., 2015). These pairwise metrics are typically applied to estimate FC among all brain regions, *i.e.*, to obtain “all-to-all” FC connectomes (Sporns et al., 2005). Networks of inter-areal FC are often represented as graphs where brain areas constitute the *nodes* (or vertices) and observed inter-areal connections the *edges* (Bullmore and Sporns 2009; Rubinov and Sporns 2010).

FC graphs estimated from MEG/EEG sensor space data are neuroanatomically uninformative and severely confounded by signal mixing. Signal mixing has two facets: first, any focal neuronal signal is picked up by several sensors. Conversely, one sensor detects a mixture of signals from several distinct sources. Source reconstruction can be used to reduce signal mixing and, importantly, elucidate the likely neuroanatomical sources of the MEG/EEG signals (Buzsaki et al., 2012; Gross et al., 2013; Hamalainen et al., 1993; Palva and Palva 2012; Schoffelen and Gross 2009). Yet, because of ill-posed nature of the inverse problem, no source reconstruction approach can yield an unambiguous estimate of the source topography. Residual signal mixing in source space, signal leakage, is quantitatively dependent on the source-reconstruction method of choice but qualitatively characteristic to all such methods.

Because of signal leakage, FC measures exhibit two distinct types of false positive observations: *artificial interactions* (AI) and *spurious interactions* (SI) (see Box 2, (Palva and Palva 2012)). AIs arise directly from the signal mixing by one true signal being smeared to multiple sensors or sources, regardless of whether true interactions are present. SIs are “ghost” interactions caused by the leakage of the signals from two true connected nodes to their surroundings nodes that in turn become falsely connected like the truly connected nodes. AIs can be suppressed by a number of bivariate metrics that typically aim to remove linear coupling terms, and therefore removing artificial and true interactions with zero-and anti-phase-lag coupling (for a review see (Palva et al., 2017)). However, the problem of SIs is much less acknowledged and more difficult to solve because SIs stem from multivariate mixing effects. With typical distributed source modeling approaches, signal leakage causes a large number of SIs that render both the network localization and graph property estimates inaccurate (Drakesmith et al., 2015). To date, one solution has been proposed for correcting SIs in oscillation amplitude correlation estimates, which simultaneously orthogonalizes all source time series through the Löwdin procedure (Colclough et al., 2015; Colclough et al., 2016). Despite this promising advance, no solutions have yet been proposed to suppress SIs for other interaction metrics.

Here we advance a novel approach, hyperedge bundling, to alleviate the problem of SIs problem in all-to-all connectivity analyses performed with any interaction metric. Instead of correcting the mixing effects in source signals *per se*, the approach is based on a quantification of the extent of mixing between all sources, evaluation of mixing similarity among all edges, and then clustering the *raw* interaction metric edges into *hyperedge* bundles. This procedure aims to yield a hypergraph where each hyperedge represents a true interaction and its spurious reflections.

In this study, we performed a large set of connectivity simulations and realistic all-to-all MEG source space analyses, in which we estimated phase synchrony as a measure of FC with an AI-insensitive metric. We show that in simulated graphs, hyperedge bundling greatly decreases the number of false positives, *i.e.*, SIs. We illustrated how bundling can support an informative visualization of FC graphs with real MEG data. We suggest that such hypergraphs constitute accurate and unbiased representations of neuronal interactions observable in MEG/EEG source space.

## 2 Theory

This section covers general topics as follows: signal mixing in MEG/EEG, how spurious interactions (SI) arise from mixing between sources; and bundling of raw edges into hyperedges. The implementations specific to this study are described in the *Methods* section. Throughout the report, we denote a connectivity graph estimated from reconstructed source time series as raw graph *G_raw_* = (*V, E*), where brain regions are nodes *v_i_ V* and interactions between nodes are “raw” edges, *e_k_* = {(*v_i_,v_J_*)∈*E*|*v_i_,v_i_* ∈ *V*}.

### 2.1 Signal mixing results in false positive artificial (AI) and spurious interactions (SI)

Let us consider a scenario where a true phase correlation is present between two distant (unmixed) sources *V_1_* and *V_2_* (Fig 1A top). The signals from *V_1_* and *V_2_* are mixed with signals of their nearby and mutually uncorrelated neighbours *V_3_* and *V_4_*. Estimating phase FC among all four nodes with the phase-locking value (*PLV*) will reveal both the true edge *E*(*V_1_*,*V_2_*) and false positive “short-range” AIs between the nearby nodes *E*(*V_1_*,*V_3_*) and *E*(*V_2_*,*V_4_*), because *PLV* is inflated by mixing (thick gray edges, Fig 1A bottom). However, due to leakage of the signal from *V_1_* and *V_2_* to their neighbors *V_3_* and *V_4_*, false positive “long-range” SIs *E*(*V_3_,V_4_*), *E*(*V_2_,V_3_*), and *E*(*V_1_,V_4_*) will also be observed (thin dashed edges). These SIs are thus only indirectly caused by mixing and, unlike the zero-phase-lag AIs (see 2.2), SIs inherit the phase-lag of the true interaction. Mixing-insensitive bivariate metrics such as the imaginary part of *PLV* (*iPLV*) can remove AIs but do not eliminate SIs if the true coupling has non-zero phase lag.

**Fig 1.**
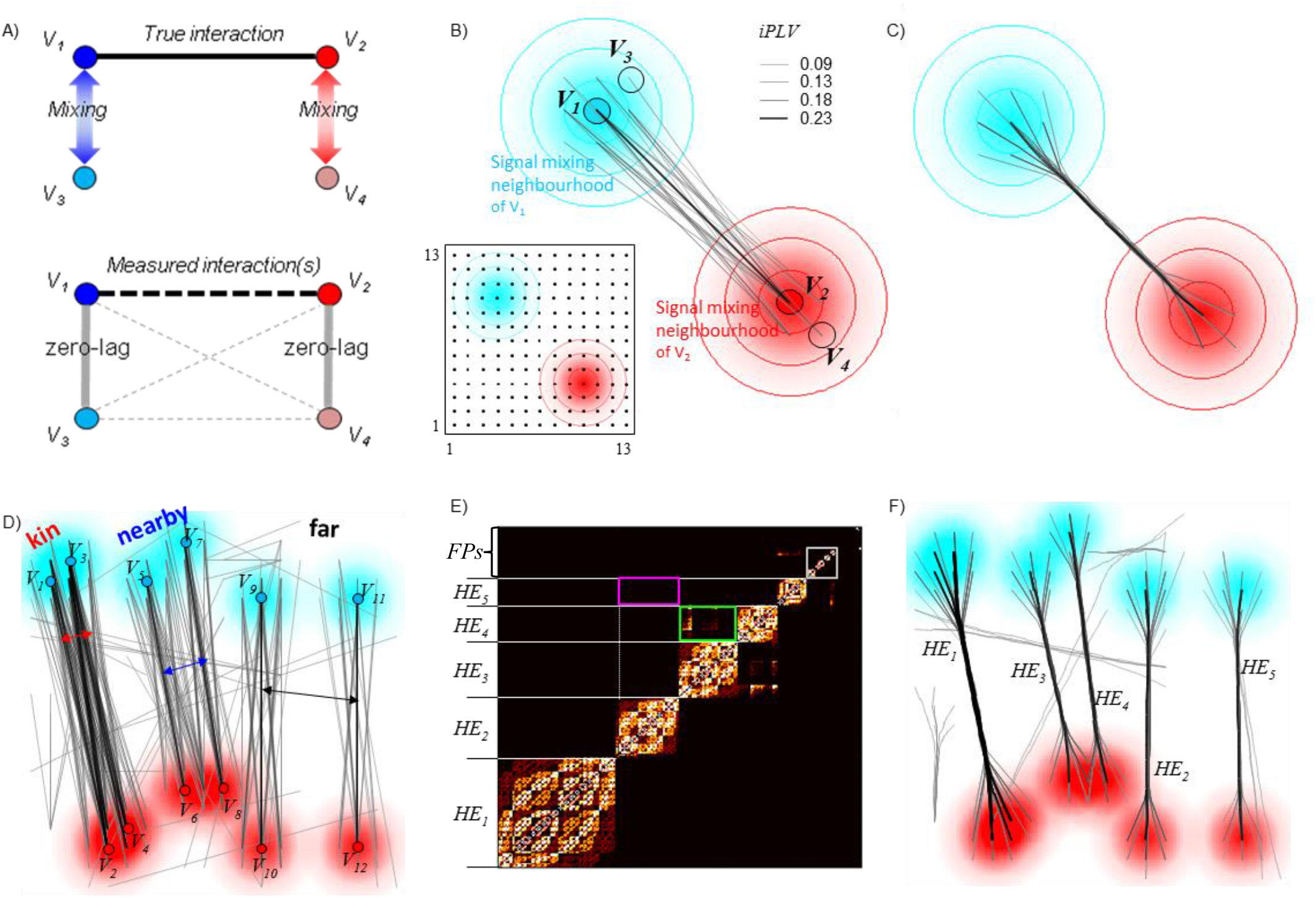
Spurious edges are indirect products of mixing and they can be bundled. ***A)*** Top: signal mixing causes the detection of artificial (AI) and spurious interactions (SI). Bottom: AIs are always zero-lag connections (solid gray edge) whereas SIs (dashed gray edges) are “ghosts” of the phase-lag of the true interaction (dashed black edge) and thus can be either zero-lag or, more often, non-zero-lag interactions. ***B)*** Toy model 1: one single true interaction *E*(*V_1_, V_2_*) on a grid of 13 x 13 point sources. Inset shows the simulated mixing neighbourhood of *V_1_* and *V_2_.* FC was estimated with *iPLV*, and the true edge (black) was discovered with multiple SIs (grey) originating from both sources’ mixing neighbourhoods. ***C***) The similarity in signal mixing between all edges (true and SI) can be quantified and all these edges can be bundled into one hyperedge. ***D)*** Toy model 2: three pairs of true edges of varying spatial distance were simulated. ***E)*** Partitioned similarity matrix *S_E_*, for toy model 2, where each row represents one edge and one cluster represents a hyperedge. The grey box indicate false-positive hyperedges; the magenta and green boxes indicate the inter-hyperedge similarity between the “far” and “nearby” pair. ***F)*** Visualization of the hyperedges defined in ***E.***

### 2.2 Quantifying the mixing between reconstructed sources

Signal mixing/leakage between two sources is instantaneous and therefore always leads to inflated zero-phase-lag correlations between the sources. Mixing does not vary over time or across frequency bands (Brookes et al., 2012; Brookes et al., 2014; Drakesmith et al., 2013; Nolte et al., 2004; Palva and Palva 2012).

#### 2.2.1 Source-reconstruction

Suppose we have a data matrix *X* = {*x_(1)_, x_(2)_*, … *x_(n)_*} ∈ ℝ^n × t^ representing narrow-band time series of *t* samples from *n* neuronal populations. Simulating a MEG/EEG recording, *X* can be linearly projected to sensor-space (Hämäläinen and Ilmoniemi 1994):

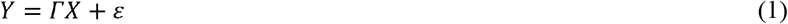

where *Y* ∈ ℝ^s×t^ represents the forward-modeled time series from *s* sensors (*n* > *s*). Here, *Γ* ∈ ℝ^s×n^ is the forward operator (or the lead field) and *ε* ∈ ℝ^s×t^ is the model prediction error derived from measurement noise. Next, *Y* can be projected back into the source-space, *e.g.*, by minimum-norm estimation (MNE) based inverse modeling:

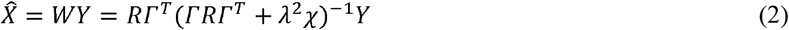

where *W* ∈ ℝ^n × s^ is the inverse operator (sources × sensors), the regularization parameter *λ^2^ = 0.1, R* is the source covariance matrix, and χ is the noise covariance matrix. Next, several thousand of source vertices can be collapsed onto a smaller number (50-400) of cortical parcels.

#### 2.2.2 Cross-talk function and resolution matrix

In MEG/EEG source connectivity studies, a resolution matrix *P* = *WΓ*(*P* ∈ ℝ^n×n^) is often used to describe the relationship between true signals and modeled signals from *n* sources in the absence of noise (Farahibozorg et al.,2017; Hauk and Stenroos 2014; Hauk et al., 2011; Liu et al., 2002). In *P*, each diagonal element quantifies the sensitivity for estimating signals from that source. Each row of *P* is the “cross-talk” function (CTF) that describes the amount of mixing between one source and all other sources. Each column of *P* is a “point-spread” function (PSFs) that describes how the modeled signal from any one source is spread across all other sources.

#### 2.2.3 The mixing function

For collapsed cortical parcels, we approximate the resolution matrix *P* with a mixing matrix *A_mix_* of dimension *n × n* parcels. Each element of *A_mix_* is a *mixing function* (*f_mix_*) that characterizes the signal mixing between two parcels. We rationalize that if the true source signals are uncorrelated, the amount of correlation at zero-lag between reconstructed signals can only be explained by mixing between the sources. Thus, *f_mix_* can be quantified by the zero-lag correlation between parcel time series estimated using a simulated MEG/EEG measurement of uncorrelated source noise.

We first generate uncorrelated signals *X_0_* ∈ ℝ^n×t^, *t* samples for *n* parcels, and forward transform them to obtain sensor signals *Y_0_* (eq. 1). We next inverse transform *Y_0_* to obtain reconstructed signals 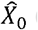 (eq. 2). In this process, the reconstructed signals 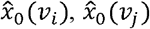 of any two nearby sources *v_i_* and *v_j_* become correlated to a certain degree due to mixing. Thus, the mixing from the simulated *“true”* signal *x_0_*(*v_i_*) to the *reconstructed* signal 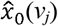 can be quantified as:

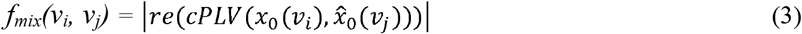

where *re()* denotes the real part of a complex number and *cPLV* is the complex-valued phase locking value (Lachaux et al., 1999):

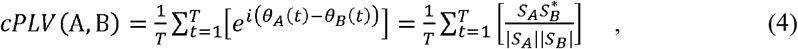

where *T* denotes the number of samples, *θ_A_* and *θ_B_* are the instantaneous phases of signal *A* and *B; S_A_* and *S_A_* are complex-valued narrow-band signals from A and B, and *z*^*^ is the complex conjugate of *z.* Because mixing is instantaneous, re(*cPLV(A,B)*) captures all correlations caused by mixing. For parcel pairs that do not become correlated by signal mixing, *f_mix_* is near zero. For parcel pairs influenced by signal mixing, *f_mix_* ≫ 0 and reaches 1 for complete mixing.

### 2.3 Signal mixing smears a true interaction into multiple spurious interactions

For a simplified illustration of how signal mixing / source leakage produces SIs, we used toy model with a 13 x 13 grid of point sources. The infidelity matrix *A_infid_* of dimension 169 x 169, was defined so that mixing between any two sources was a 2D Gaussian distribution decreasing with distance between the two sources (inset, Fig 1B, methods see *Supplementary*).

We simulated one true edge by setting two sources *V_1_* and *V_2_* to have phase coupling of 0.9 with nonzero phase lag and keeping the remaining 167 sources uncorrelated. Next, we introduced mixing between reconstructed sources and mapped all-to-all phase FC with an AI-free metric, the imaginary part of the phase-locking-value (*iPLV*) (Palva and Palva 2012)

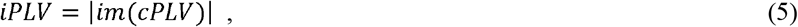

The *iPLV*, like the imaginary coherency (Nolte et al., 2004), removes zero-lag couplings by excluding the real part of *cPLV.* Therefore, *iPLV* yields only the true phase-lagged interactions and their false positive ghosts (SIs). In this simulation, visualization of the strongest 0.1% of *iPLV* edges revealed the true edge and several SIs, all of which connected sources within the mixing neighbourhoods of the true sources *V_1_* and *V_2_* (Fig 1B).

### 2.4 Raw edges can be bundled into hyperedge by their mixing similarity (*S_E_*)

The *mixing similarity* can next be derived with the known mixing matrix *A_mix_* to describe how close these edges are with each other in signal mixing. A bivariate similarity estimation yields a mixing similarity matrix *S_E_*, where each element *S_E_*(*i, j*) quantifies the similarity between two edges *E_i_*, *E_j_* (for how-to, see 2.5).

Our objective is to classify raw edges by mixing similarity into “hyperedges”, where each *hyperedge* is a “bundle” of raw edges (including true and false-positive SI edges): *HE_κ_* = {{*e_k_* = (*v_i_,v_j_*)} ∈ *E*/*v_j_,v_i_* ∈ *V*}. The raw graph is thereby transformed into a hypergraph *G_h_* = (*V*, *HE*). Within any one hyperedge, all raw edges are mixing-wise close to each other but distant from the raw edges of other hyperedges, and thus collectively representing a “community” of raw edges that we hypothesize to include the underlying true interaction and its ghosting SIs.

This classification can be done by partitioning the *S_E_* matrix into clusters with an appropriate clustering method. In the toy model, bundling transformed the raw graph with a multitude of false positives into a hypergraph with one hyperedge that captured the true interaction with zero false positives (Fig 1C).

For visualizing hyperedges, we utilized a “force directed edge bundling” method that both indicates the adjacency of the constituent raw edges and illustrates the loci where the SIs originated (Holten and Wijk 2009).

### 2.5 Hyperedge bundling for multiple true interactions

To demonstrate that bundling could be extended to separate multiple true interactions, we expanded the simulation and modeled interactions with three degrees of adjacency: “kin”, “nearby”, and “far”. The estimated raw graph yielded the true-positive (TP) edges surrounded by numerous false positive (FP) SIs (Fig 2D). Estimating and partitioning the edge similarity matrix *S_E_* revealed that: 1) two “kin” edges were inseparable and together with their SIs they merged into the largest hyperedge *HE_1_* (Fig 2E); 2) the “far” pair was clustered into two clearly separable hyperedges *HE_2_* and *HE_5_*; 3) the “nearby” pair and their SIs were also clustered into two distinct hyperedges *HE_3_* and *HE_4_* with greater inter-hyperedge similarity as measured by mean-linkage (green box) than the “far” pair (magenta box); 4) a few scattered random false positive edges were also clustered into hyperedges (gray box), but they were much smaller in size than any of the hyperedges containing a true edge.

**Fig 2.**
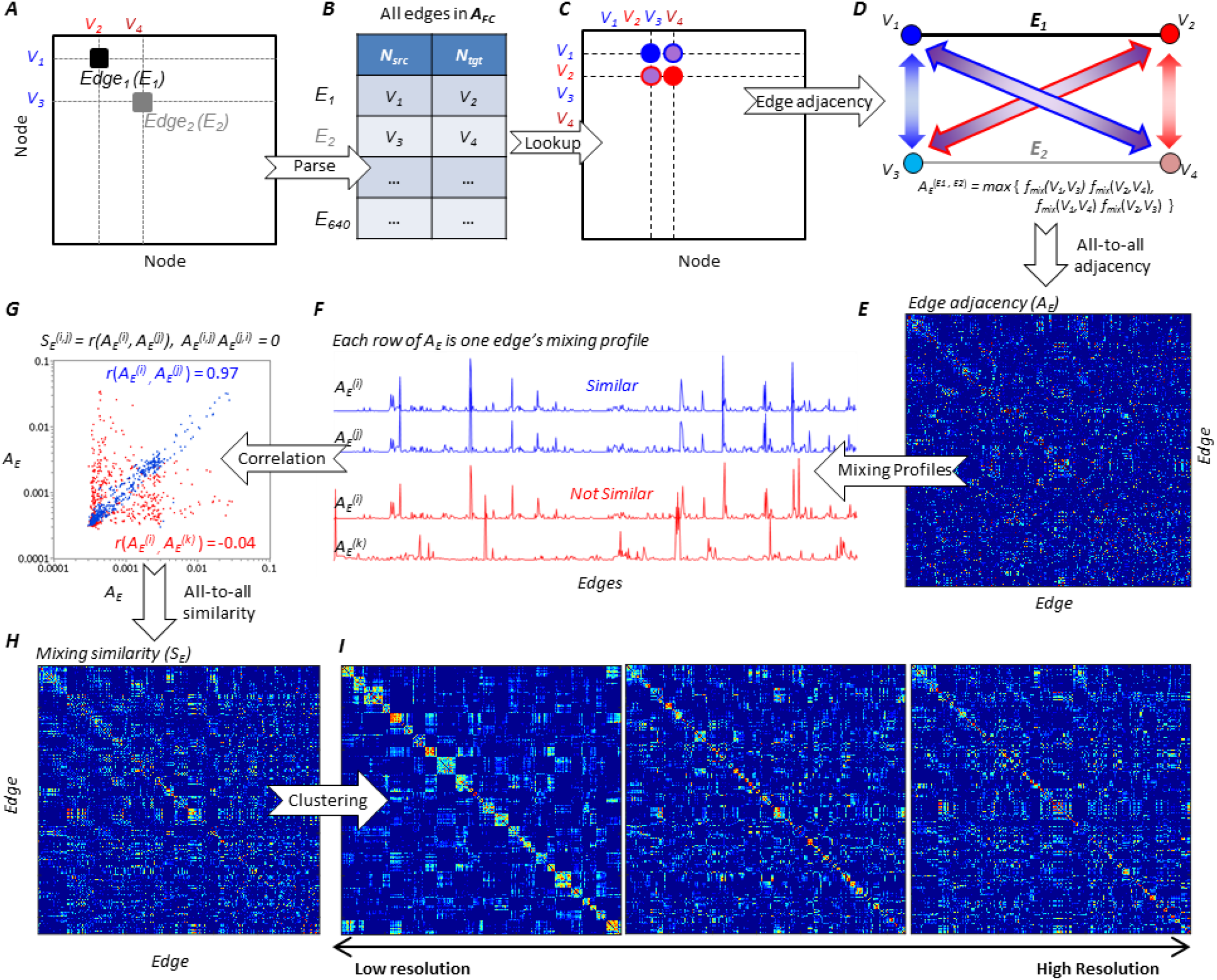
Bundling of raw edges into hyperedges. ***A)*** The true interaction *E_i_* and one of its SIs *E_2_* from Fig 1B schematically shown in matrix form. ***B)*** The raw graph *A_FC_* (a sparse matrix containing only significant edges) is parsed to a list node pairs, each pair representing one edge. ***C)*** For *E_1_* and *E_2_*, the mixing (*f_mix_*) between all of their constituent nodes can be found in the mixing maxtrix *A_mix_.* ***D)*** The edge adjacency (*A_E_*) between *E_1_* and *E_2_* is the maximum product of constituent nodes’ mixing. ***E)*** *A_E_* is computed for all the pairs of edges found in *A_FC_.* Data taken from a randomly selected simulation. ***F)*** Examples of edges that are similar (blue) and not similar (red) in their mixing profiles. ***G)*** Similarity between two edges is the correlation between two edges’ mixing profiles. ***H)*** Mixing similarity matrix *S_E_*. ***I)*** The partitioning of this *S_E_* at low, medium and high resolutions.

If a hyperedge containing at least one true raw edge is considered as a TP observation, bundling greatly decreased graph noise in terms of the FP/TP ratio. FP/TP in raw graph was 239/6 and 4/5 in the hypergraph, which marks a reduction in the fraction of FPs by a factor of 50. Visualizing these bundles showed that the hypergraph had less visual clutter and facilitated identification of the true interactions compared to the raw graph (Fig 2F).

### 2.6 Estimation of the edge similarity matrix *S_E_*

Hyperedge bundling is based on the raw connectivity graph *A_FC_* (a sparse matrix containing only significant edges), and the mixing matrix *A_mix_* (Fig 2A, C). We first parsed the edges in *A_FC_* into a list of node pairs (Fig 2B). We next find the mixing function *f_mix_* between all involved nodes from *A_mix_* (Fig 2C, and illustrated geometrically in Fig 2D) to compute the edge-to-edge adjacency in signal mixing.

#### 2.6.1 The edge adjacency matrix (*A_E_*)

For a raw graph with *m* edges, the edge-to-edge adjacency matrix *A_E_* ∈ ℝ^*m × m*^ represents the pairwise mixing adjacency among all raw edges and is necessary for computing the similarity matrix *S_E_.* The adjacency between two edges *E_i_*(*V_1_*,*V_2_*) and *E_j_*(*V_3_,V_4_*)} was defined as follows (Fig 2D):

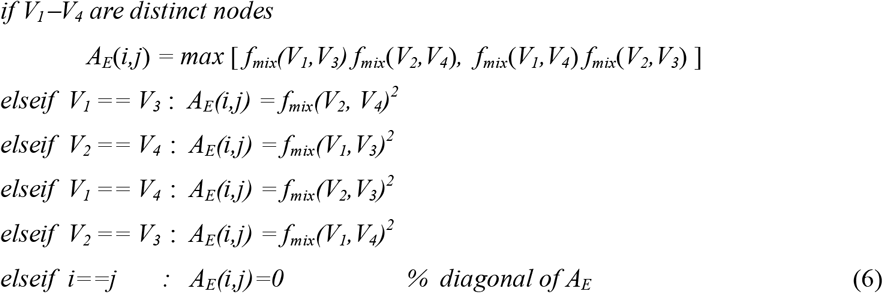

here “==“ is assertion, “=“ is assignment. This algorithm is applied for all pairs of edges in the raw graph to populate the *A_E_* matrix (Fig 2E).

#### 2.6.2 Evaluation of Edge Similarity (S_E_) with correlation of edge mixing profiles in A_E_

We denote rows of the *A_E_* matrix as the *signal mixing profiles* so that *A_E_(i)* and *A_E_(j)* are the mixing profiles of edges *E_i_* and *E_j_*, respectively, and thus indicate their mixing adjacency to all the other raw edges in the graph. If *E_i_* and *E_j_* are similar to each other, *i.e.*, a high correlation between *A_E_(i)* and *A_E_(j)*, edge *E_i_* will be similar to all the edges in the raw graph that *E_j_* is similar to, and vice versa (Fig 2F&2G). Such pattern can be already observed in the simplified models (Fig 1) where SIs of any given true edge are all close to each other and adjacent to the true interaction.

Conversely, if two edges are far apart in mixing, their mixing profiles exhibit little to no correlation. Using correlation estimates of mixing profiles, it is thus possible to assess the significant similarity of all pairs of edges in *A_E_* and populate the similarity matrix *S_E_* ∈ ℝ^*m × m*^ (Fig 2H). Hyperedge bundling is based on the notion that a *S_E_* can be partitioned into clusters of raw edges that are similar to each other in mixing within each cluster and therefore to collectively reflect a shared true underlying interaction.

### 2.7 The resolution of hyperedge bundling is defined by the cutoff limit

We partition the edge similarity matrix *S_E_* into clusters of “hyperedges” so that within any one hyperedge, the raw edges are mixing-wise close (large *S_E_* values) to each other and distant (small *S_E_* values) from raw edges of other hyperedges.

We now introduce a control parameter, the *cutoff limit* (CL) that dictates the “resolution” of a hypergraph. CL is defined as the ratio of desired number of clusters to the number of available raw edges to be clustered. For example, for a graph of 1000 edges, a CL of 0.1 causes the clustering method to partition the *S_E_* matrix into 100 hyperedges. We chose to control clustering using the CL for better comparability of clustering methods or graphs of different sizes. The similarity matrix *S_E_* ∈^*m* × *m*^ can be partitioned into arbitrary number of clusters from 1 to *m* - 1, *i.e.*, CL ranging from 1/*m* to (*m*-1)/*m* (Fig 2I, for technical details on how CL is related to the depth at which dendrogram was cut into clusters, see *Supplementary*).

### 2.8 Validate the stability of hyperedge clustering

To ensure that the hyperedges are not random outcomes of partitioning the similarity matrix, the “stability” of partitioning solutions must be evaluated. We ask, at any resolution (CL=c), if the differences between the partitioning solutions of *n* randomly perturbed versions of a similarity matrix *S_E_* is statistically smaller than their surrogate counterparts, the partitioning solution can be considered as stable *(Supplementary).* The distance between two partitioning solution can be estimated with the *variation of information* (VI, (Meilă 2007)). The independent perturbations to a similarity matrix can be acquired by randomly deleting a small subset, *e.g.*, 10 or 20%, of the elements in the similarity matrix (Ben-Hur et al., 2002; Williams et al., 2015). The surrogates can be obtained by randomly permuting the perturbed similarity matrix.

## 3 Methods

The goal of this study was to assess the performance and applicability of hyperedge bundling in suppressing spurious interactions (SI) in MEG/EEG source connectivity studies. To this end, we obtained large numbers of functional connectivity (FC) graph estimates from simulated data with realistic sources and inverse modeling. We next evaluated the efficacy of hyperedge bundling in capturing true positive (TP) interactions and rejecting false positive (FP) SIs. Finally, we demonstrated the bundling of FC graphs estimated from MEG data recorded in a visual working memory (VWM) experiment.

This section includes the procedural outlines of the simulations and evaluation of bundling efficacy. The preprocessing pipeline, technical details of the simulations and preprocessing of the VWM experiment are described in *Supplementary.* The Python 2.7 and National Instruments ^TM^ LabVIEW version of the hyperedge bundling program can be downloaded from: https://figshare.com/projects/Hyperedge_Bundling/26503.

### 3.1 Simulating “truth” time series of varying coupling strengths

In real electrophysiological data, mixing across source loci and subjects is inhomogeneous (Brookes et al., 2014) and coupling strengths of neuronal interactions also exhibit great spatiotemporal and inter-subject variability (Preti et al., 2016; Zalesky et al., 2014). To account for such variability, we created 1000 distinct *truth* graphs each containing 200 randomly generated true interactions between 400 cortical parcels in a standard cortical source space (Destrieux et al., 2010). Each node thus connected only to a single other node, which allows an unbiased survey of the whole cortical surface in every graph realization. We did not simulate structured networks therefore excluding the impact of higher order SI. These higher order SI can arise from common drive, third-party sources, and cascade effects, although identifying them is of equal importance (Marinino and Bressler 2015; Wollstadt et al., 2015).

For every truth graph, we simulated ten sets of coupled time series, representing two different modes of coupling, *i.e.*, gamma distribution (*C_λ_* with maximum coupling of 0.9 and order parameter *r* ranging from 1 to 20) or uniform distribution (*Cc*) at 5 different levels of coupling strength each (*Supplementary*). A set of uncorrelated null hypothesis time series was also simulated for each truth graph. These null hypothesis time series were used for estimating the parcel mixing properties (3.2) and as the baseline condition against coupled conditions in group analysis.

### 3.2 Estimation of mixing properties using the *H_0_* time series

Mixing in source reconstructed MEG/EEG data is essentially captured in the forward and inverse operators used in source reconstruction. These operators are determined by the data acquisition system and specifics of the individual source model (Wens 2015). In addition to the mixing function *f_mix_* (see 2.2.3), we characterized the source model used here with a set of additional mixing metrics obtained from the 12 subjects from the VWM experiment:

1) Parcel fidelity quantifies the reconstruction accuracy and is defined as the phase correlation between the *simulated* null hypothesis time series *x*_0_, and *reconstructed* null hypothesis time series 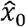 of parcel *v_i_*

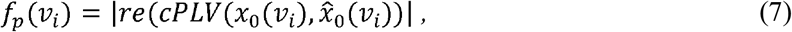
2) Edge fidelity, *f_e_*(*v_i_,v_j_*) = *f_p_*(*v_i_*)*f_p_*(*v_j_*), that quantifies the reconstruction accuracy of raw edges connecting two parcels *v_i_* and *v_j_.*
3) Residual spread function is the correlation between two parcels reconstructed null hypothesis time series.

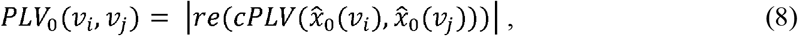

The definition of *PLV_0_* appears similar to that of *f_mix_*, but they are conceptually different. The *f_mix_* measures how much of each source’s true signals are picked up in other sources’ reconstructed signals. *PLV_0_*, on the other hand, is the correlation between any two sources’ modeled time series that both are contaminated by mixing with numerous other sources. Because the *iPLV* estimates can be biased by mixing, we used *PLV_0_* to exclude edges connecting sources with large mixing (Palva et al., 2017).

### 3.3 Elimination of poorly measurable edges with the intractable-edge-mask (IEM)

We applied an intractable-edge-mask (*IEM*) to exclude edges that connect sources with poor reconstruction accuracy. True interactions between these sources may exist, but cannot be reliably detected because estimations of connectivity between them are unreliable due to the limitations of the source model. We utilized the mixing properties (see 3.2) and construct a group-level *IEM* in two steps:

1. With average edge-fidelity <*f_e_* and the residual spread <*PLV_0_* >, we create two Boolean masks:
  i. The edge-fidelity mask (*M_fe_*) to exclude edges with low fidelity, thereby removing edges connecting poorly reconstructed sources.
  ii. The residual spread mask (*M_PLV0_*) to exclude edges with large *PLV_0_*, thereby removing edges whose FC estimates likely are much distorted by mixing between these loci (Palva et al., 2017).
2. The *IEM* is the union of these two masks.

In this study, we set 0.1 as the threshold for *M_fe_*, which removed the 40% most poorly reconstructed edges from all 79,800 (*N(N—1)/2*, *N* = 400) possible edges in raw graphs. The *M_PLV0_* was acquired by deleting edges whose *PLV_0_* was greater than the 95^th^ percentile of the *PL V_0_* matrix.

### 3.4 Estimation of group-level FC of simulated graphs

The group-level significant *iPLV* estimates thresholded with the *IEM* were used as raw graphs for hyperedge bundling. The group-level analysis for the simulated graphs and for real MEG/EEG data in the VWM experiment were carried out in the same manner. For simulated graphs, we forward-and inverse-modeled the coupled truth time series into 12 subjects’ individual source space, thereby introducing mixing into reconstructed signals (Schoffelen and Gross 2009). We next estimated *iPLV* connectivity for these subjects. We then tested across subjects, for each edge in every estimated FC graph, whether there was a significant difference (one-tailed t-test) in the *iPLV* estimate between the coupled and the *H_0_* condition. Those edges that showed a significant difference were identified as raw edges (corrected for multiple comparisons within each FC graph). We acquired FC graphs with three significance levels p < 0.05, 0.01, and 0.001 for each of the ten coupled time series.

### 3.5 Hyperedge bundling with two clustering methods

After applying the *IEM* to all group-level FC matrices, we followed the procedures described in *Theory* to obtain the similarity matrix *S_E_* for each FC. We next partitioned each *S_E_* into clusters of “hyperedges” with two clustering methods. The unweighted pair group method with arithmetic mean (UPGMA) is an agglomerative hierarchical clustering method that builds a rooted hierarchical tree to represent the distance in signal mixing between all raw edges (Jain et al., 1999). The Louvain method for community detection extracts communities by optimizing the modularity of clusters through a gradient descent procedure (Blondel et al., 2008).

### 3.6 Comparing hypergraphs with raw graphs

We denoted the TPs as the edges from truth graphs that were identified as significant edges in the group-level FC matrix, and FPs as significant edges in the group-level FC matrix but absent in the truth graph. Thus, the true positive rate (TPR, sensitivity) is given by *TPR* = *TP/N_true*_*, where *N_true*_* is the number of “detectable true edges” referring to the number of simulated true edge that passed the intractable-edge-mask. We further defined the *noise* as the FP to TP ratio. An ideal group-level FC should capture as many of the true interactions as possible while rejecting other edges, *i.e.*, high TPR and low FP/TP.

We used TPR and FP/TP as the main criteria to characterize raw graphs instead of the commonly used receiver operating characteristic curve (ROC) for two reasons. First, the ROC is derived from the TPR and false positive rate (FPR) which are not directly comparable between raw graphs and hypergraphs, as these are different constructs; second, because the number of FP is disproportionally larger than that of TP (as shown later with an example), the shape of the ROC is misleadingly optimal when limiting the number of raw edges

We defined a TP hyperedge (TP_HE_) as a hyperedge capturing at least one TP raw edge, whereas a FP hyperedge (FP_HE_) contained only FP raw edges. Hyperedges may also contain multiple TP raw edges. To quantify this, we defined *separability* as the fraction of true positive hyperedges that contain only one TP raw edge out of all true positive hyperedges. An ideal hypergraph should balance high TPR and separability against low FP/TP.

## 4 Results

This section includes three parts: 1) Demographics of group-level FC of the simulated graphs; 2) Efficacy of hyperedge bundling; 3) Application of hyperedge bundling to real MEG data.

### 4.1 Group-level FC as raw graphs

In individual subjects, mixing introduced by the virtual MEG experiment distorted *PLV, iPLV* and the phase-lag of all measured graphs of varying coupling strength including the *H_0_* time series (*Supplementary*). To find group-level significant edges, we tested for each edge whether there was a difference in *iPLV* value between the coupled condition and the *H_0_* condition (Fig 3A, see 3.4). Edges that showed a significant difference were reported as raw edges (corrected for multiple comparisons). Thus, we obtained FC graphs for each of the ten sets of coupled graphs at 3 significance levels of *p* < 0.05, 0.01 and 0.001.

**Fig 3.**
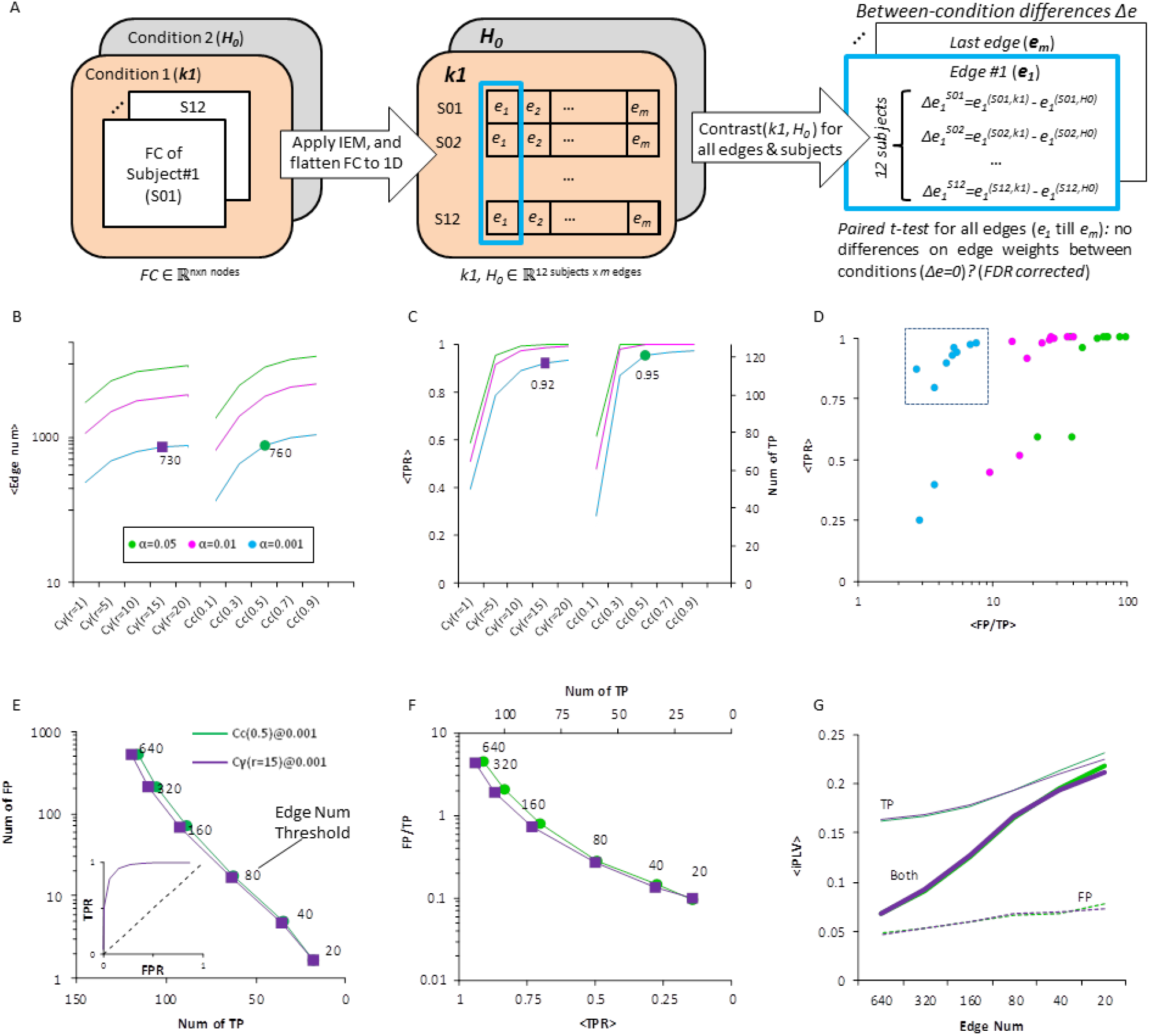
The demographics of group-level FC of simulated graphs. ***A)*** Significant edges were determined with a paired one-tailed t-test between a coupled-edge condition (k1) and the *H_0_* condition for simulated graphs. ***B)*** For initial evaluation of bundling, we chose one set of gamma-distribution-coupled (C_λ_) and one set of uniform-distribution-coupled (Cc) graphs, which are indicated by the markers. *C)* True positive rate TPR (see methods) of the two chosen graphs was above 90%. ***D)*** The true positive rate (TRP) as a function of noise (FP/TP) for all coupling strengths. ***E)*** In the chosen sets of graphs, the number of FP decreases exponentially, while the number of TP decreases linearly. Inset shows the ROC of C_γ_ edge weights threshold. ***F)*** Noise (FP/TP ratio) as a function of TPR. ***G)*** The mean *iPLV* of TP or FP edges alone, and all edges.

#### 4.1.1 Raw graphs of iPLV edges are noisy

Overall, the number of significant *iPLV* edges increased as coupling strengths increased (Fig 3B). The group-level graphs at all 3 significance levels captured over 75% of all detectable TP edges, except in the case of weak uniform coupling, Cc(0.1) (Fig 3C). We simulated 200 random edges in each ground truth graph and computed the true positive rate (TPR) for each measured group graph as the number of significant edges divided by the number of all simulated true edges that passed through the intractable-edge-mask (IEM). Despite the high TPR, there was a large variability in the ratio of false and true positives, FP/TP, across these graphs (Fig 3D).

#### 4.1.2 Is strict statistical thresholding a good solution for pruning FPs?

We chose the graphs of gamma coupling (*C_λ_*) with order parameter *r* of *15* and uniform coupling (*C_C_*) with coupling *of 0.5* to test statistical thresholding (below) and hyperedge bundling because they had comparable TPR (Fig 3C) and equivalent true edge strengths (see distribution in Supplementary 1). Moreover, both contained only ∼750 edges, which mitigated computational overhead in later clustering analyses.

One sensible way to identify key structures in FC graphs is to apply a statistical threshold to *iPLV* values. We found that by increasing the significance *iPLV* threshold, the number of FP edges decreased at a faster rate than the number of TP edges in both graphs (Fig 3E). Around 120 of the 640 strongest edges were TP, giving a TPR > 90% for 125 detectable true edges, but a FP/TP ratio of 4. When retaining the 20 strongest edges reduced the FP/TP to 0.1 (Fig 3F) but at the cost of reduced TPR, (TPR = 0.15). Overall we found that the mean *iPLV* of TP edges was larger than that of FP edges’ (Fig 3G), which suggests that strict thresholding is an applicable solution for reducing FP/TP but comes at a price of an elevated false negative rate, although the shape of ROC curve appeared to be optimal (inset Fig 3E).

### 4.2 Hypergraphs yields better FP/TP than raw graphs with reasonable TPR cost

#### 4.2.1 The stability of clusters

Evaluating the stability of clustering was a necessary step prior to further analysis of the properties of hyperedge clusters. The resolution of clustering and thereby of the hypergraphs was controlled by the *cutoff limit* (CL, see 2.6). We used bootstrapping to identify the CL range that yielded stable partitioning of the raw graphs (see Methods and *Supplementary).* We found that at CL < 0.4, both UPGMA and Louvain clustering yielded significantly more stable partitions for simulated graphs than their randomly rewired counterparts (Fig 4A). For the 640 raw edge graphs, this CL upper bound corresponded to ∼250 hyperedges. In the following analysis, we thus tested bundling with CL ranging from 0.05 to 0.45.

**Fig 4.**
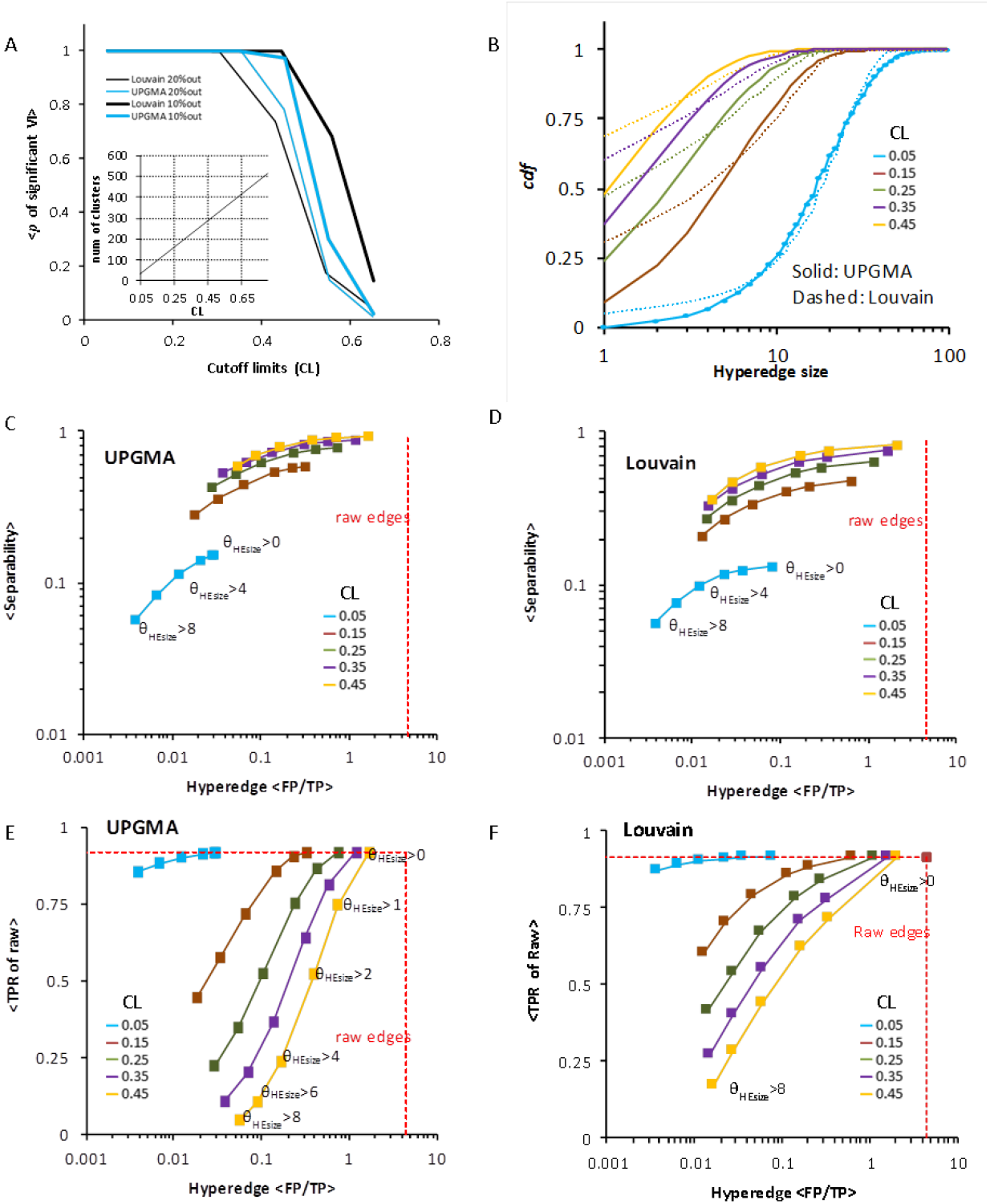
Hyperedge bundling outperformed raw edges. ***A)*** The hypergraphs created with both clustering methods were stable below CL of 0.4. ***B)*** The cumulative distribution function (cdf) of hyperedge size at different levels of CL, computed with hyperedges pooled from 500 graphs with 100 iterations within each graph. For both clustering methods *C)* and *D)*, increasingly strict hyperedge size threshold (*θ_HEsize_* varying from 0 to 8) caused separability and noise level (FP/TP) to decrease. ***E,F***) The retained true positive raw edges also decreased as hyperedge size threshold increased.

#### 4.2.2 Cluster-size distribution

We next quantified the distributions of hyperedge sizes (numbers of raw edges per hyperedge, Fig 4B) by pooling hyperedges from 500 clustered graphs with CL ranging from 0.05 to 0.45. As expected, we found a systematic shift towards smaller hyperedges with increasing resolution/CL. The Louvain method consistently yielded more small hyperedges than UPGMA.

#### 4.2.3 Hyperedge-bundling performance: trade-offs between separability, TPR and graph noise

Hyperedge bundling aims to detect and separate as many TP interactions as possible while rejecting as many FP as possible. We tried to find an optimal balance among these competing outcomes by taking into account two aspects of hyperedge bundling: separability and noise. We defined *separability* as the ratio between singleton TP hyperedges (containing only one TP raw edge) and all TP hyperedges, and *noise* as the FP/TP ratio of the hyperedges. An ideal hyperedge partitioning would thus have *separability* = 1, FP/TP ∼0, and a TPR equal to the TPR of raw edges.

We observed that by increasing the hyperedge resolution (CL from 0.05 to 0.45), the *separability* increased but noise also increased with both clustering methods (Fig 4C, 4D). Thus at coarse resolutions (low CL), multiple TP raw edges were partitioned into one hyperedge but there were very few FP hyperedges, likely because there were less small-sized hyperedges. Conversely, at fine resolutions (high CL), separability was improved but at the cost of having greater numbers of FPs.

Knowing that small hyperedges are more likely to be FPs than large hyperedges (Fig 1E), we further tested whether excluding hyperedges by size would decrease noise. At each resolution level, excluding small hyperedges lead to a decrease in noise (FP/TP decreased with increasing *θ_HEsize_*, Fig 4C, D). Nevertheless, this was accompanied by reduced separability (y axis, Fig 4C, D) and a reduced TPR (Fig 4E, F) caused by the removal of small-sized TP hyperedges together with FP hyperedges.

To summarize, at all graph resolutions, hypergraphs were less noisy than raw edge graphs. In the least noisy hypergraph (*e.g.*, Louvain, CL = 0.05 and *θ_HEsize_* > 8), 87% of the 125 TP raw edges were retained while achieving a 10^3^-fold decrease in noise compared to the underlying raw graphs, *i.e.*, FP/TP decreased from (640-125)/125 = 4.1 (C_γ_ raw graphs in Fig3E) to 3.8×10^-3^ (leftmost filled box on the cyan curve, Fig 4F). Nevertheless, this improvement came at the cost of poor separability, meaning many hyperedges in CL = 0.05 graphs contained several true edges. To balance an optimal trade-off, we decided to use CL ≥ 0.15 and *θ_HEsize_* ≥ 2, expecting to achieve a reduction of FP/TP to 0.1 (from 4.1 in raw edges) with negligible reduction in TPR and adequate separability (0.5).

#### 4.2.4 Louvain clustering yields less noisy hypergraphs but lower separability than UPGMA clustering

The Louvain method produced more small hyperedges than the UPGMA method (Fig 4B). Although the Louvain hypergraphs had higher level of noise when retaining singleton hyperedges (*θ_HEsize_* = 0), this relation was inverted when singleton hyperedges were screened (Fig 5A). This indicates that the majority of the singleton hyperedges yielded by Louvain were FPs. Moreover, the Louvain hypergraphs had greater TPR when CL values were between 0.15 and 0.25 (Fig 5B). These advantages, however, came at the cost of separability, which was better with UPGMA throughout the tested range (Fig 5C).

**Fig 5.**
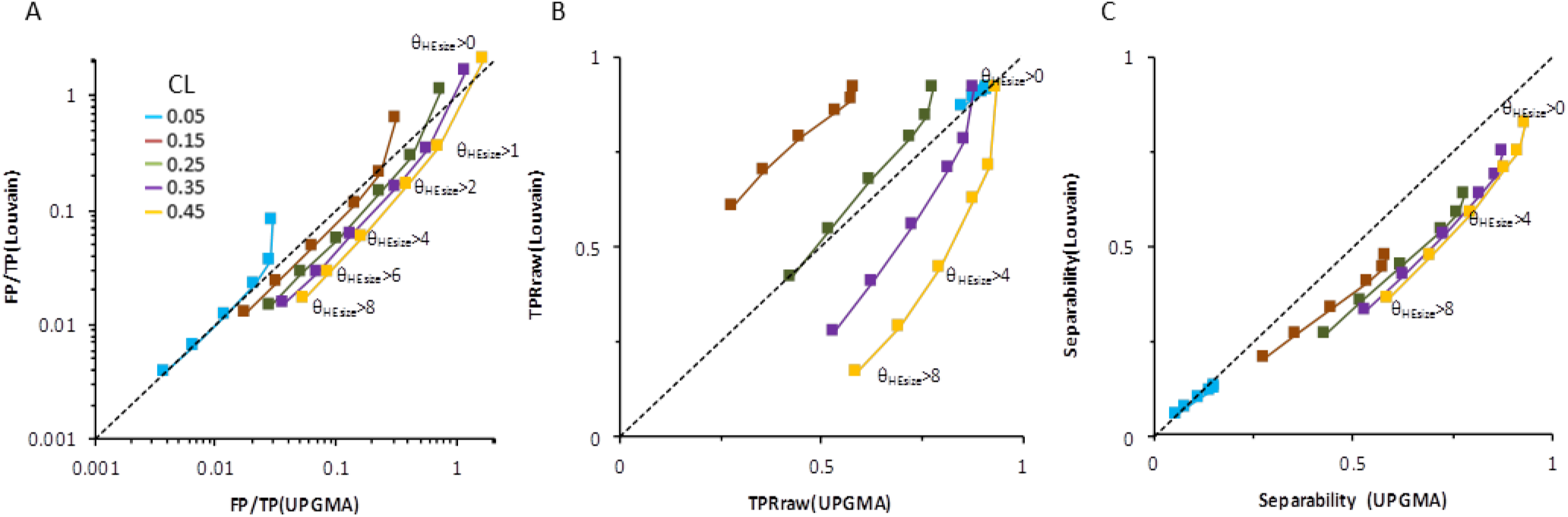
Louvain clustering method yielded hypergraphs with lower noise but also lower separability than UPGMA. *A)* For CL values 0.15 - 0.45, Louvain hypergraphs had lower noise after singleton hyperedges were deleted. *B)* True positive rate TPR was larger in Louvain hypergraphs for CL values 0.15 and 0.25 and larger in UPGMA hypergraphs for CL values 0.35 and 0.45. *C)* Separability was higher for UPGMA method.

### 4.3 Visual working memory networks: real MEG data

To assess the feasibility of using hyperedge bundling with real MEG/EEG data, we applied bundling to raw graphs that reflected significant strengthening of inter-areal phase synchronization during memory retention compared to pre-stimulus baseline during a visual working memory task (see *Supplementary* and Honkanen et al., 2015).

We found that the *iPLV* estimates in alpha-and gamma-frequency band were greater during memory retention than in pre-stimulus baseline. Here, we picked the 1000 strongest *iPLV* edges and drew them as lines linking the synchronized parcels on a flattened cortical surface (Fig 6A, 6B). We also illustrated a randomly picked graph from our simulations as a comparison (Fig 6C). We applied hyperedge bundling (UPGMA with CL = 0.15, *θ_HEsize_* > 6) to these raw graphs. The resulting hypergraphs, the real MEG and simulated FC graphs alike, offer better visualization of large-scale FC than raw graphs, emphasizing the long-range synchronizations between brain regions(Fig 6D, 6E, 6F).

**Fig 6.**
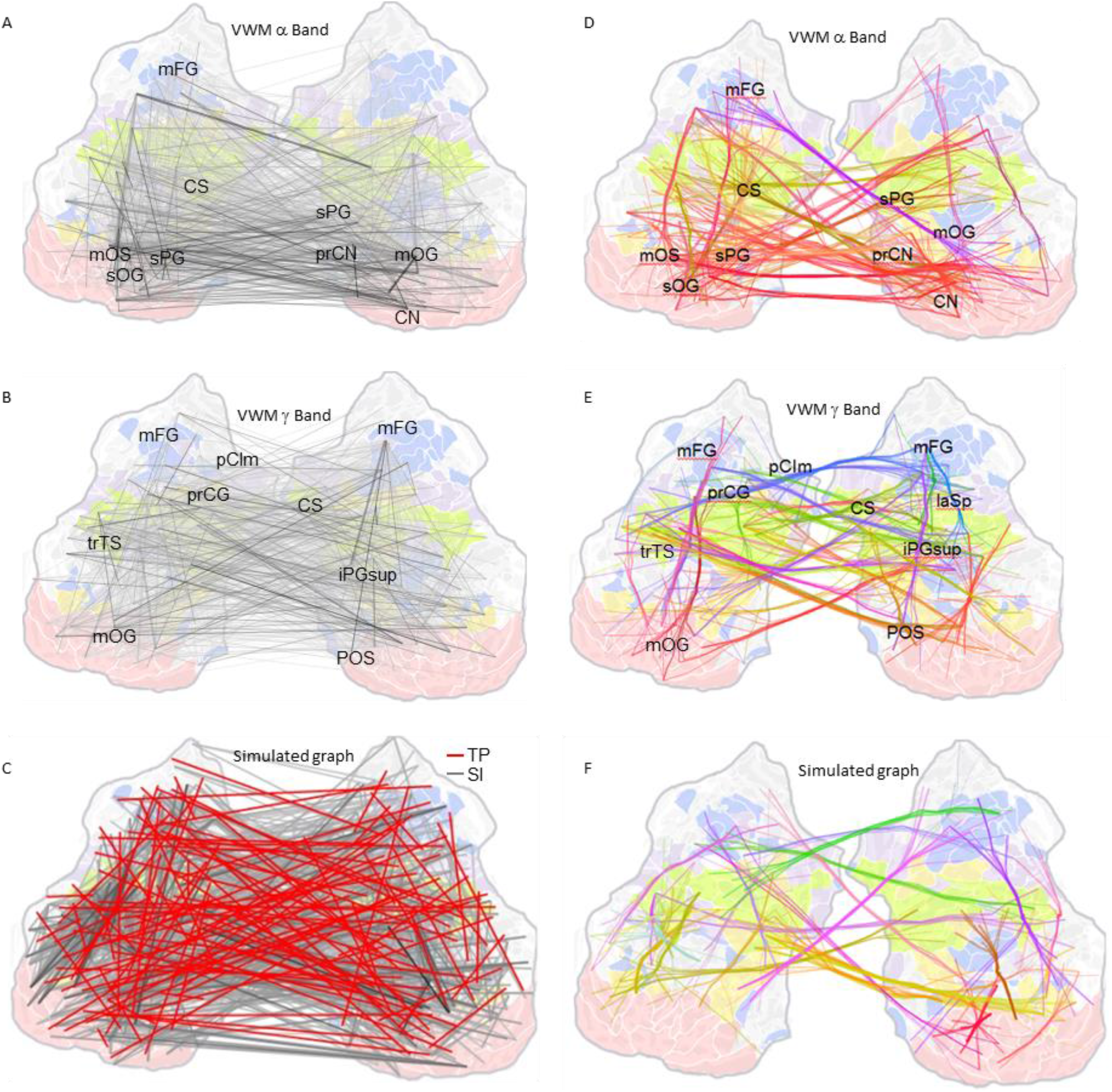
Hypergraphs improve visualization of real and simulated data. Visual crowding of numerous group-level *iPLV* edges of 1:1 phase synchronization in ***A)*** alpha and ***B)*** gamma frequency band during VWM retention (real MEG data), ***C)*** a simulated graph overlaid on a flattened 2D map of cortical regions. ***D, E, F)*** Hypergraphs of ***A,B,C). D)*** In alpha band, bundles of long-range hyperedges connect occipital and parietal areas. Hyperedges were created with CL=0.15, *θ_HEsize_* > 6. ***E)*** In gamma band, long-range hyperedges were observed in the frontal and central regions. On these 2D map, different parcel colours indicate functional subsystems defined by (Yeo et al. 2011) and in hypergraphs, edge colours are obtained by mixing of the colours of connected parcels. CN: cuneus; CS: central sulcus; iPGsup: supramarginal gyrus; mFG: middle frontal gyrus; mOG: middle occipital gyrus; mOS: middle occipital sulcus and lunatus sulcus; laSp: posterior ramus; prCG: precental gyrus; pCIm: middle posterior cingulate; prCN: precuneus; sPG:superior parietal lobule; sOG: superior occipital gyrus.

## 5 Discussion

MEG and EEG have great potential for yielding insight into the spatio-temporal structure of brain connectivity. Nonetheless, due to the ill-posed nature of the inverse problem, linear mixing and inaccurate source localization complicate MEG/EEG connectivity analyses both by distorting phase and amplitude estimates and by leading to false positive observations of artificial (AIs) and spurious interactions (SIs). We advance here a novel methodological framework, hyperedge bundling, to suppress SIs in brain connectivity graphs. We found that hyperedge bundling can be used to reduce the false positive rate with moderate to little decrease on the true positive rate.

Hyperedge bundling has several features that are advantageous and facilitate its application. First, since it is done only after interaction analyses, it does not require sophisticated preprocessing to suppress mixing effects in the original source time series. Hyperedge bundling only requires the forward and inverse operators and a mixing function estimated analytically or from simulations. Accordingly, it also inherently takes the source-model heterogeneity appropriately into account. Hyperedge bundling is also independent of the interaction metric and can be applied to connectomes estimated with any bivariate interaction metric. Finally, the nodal groups in the hypergraph obtained from hyperedge bundling constitute data-driven coarsening of originally high-resolution source parcellations. We suggest that these nodal groups be more representative of the true co-active local areas than *a priori* constructed low-resolution parcellations. This can be an aspect for future work.

In summary, hyperedge bundling can be used to suppress SIs and identify putative true edges in brain connectivity data and thereby to improve the localization of true interacting neuronal networks.

### Hyperedge bundling vs. edge thresholding: reducing FP/TP while maintaining acceptable true positive rate

Some connectivity studies have reduced the amount of edges by applying strict criteria on edge selection. However, biases and instability of graph properties can be introduced when using arbitrary threshold criteria on raw edges (Drakesmith et al., 2015; van Wijk, Bernadette C. M. et al., 2010) and weak connections may also play an important role in cognitive functions (Santarnecchi et al., 2014). Nevertheless, imposing strict criteria for thresholding is an attractive option for increasing the fraction of true positives among all observations, *i.e.*, decreasing the FP/TP ratio (see Fig 3E and F) and for focusing the outcome on most robust effects. However, this approach, while effective in excluding FPs (SIs), also excludes a large fraction of true positives. For example, we found that in raw graphs when we applied a threshold strict to decrease noise (FP/TP ratio dropped from 4 to 0.1), but the TPR was reduced to 0.15. In contrast, with hyperedge bundling we could obtain the same noise level (FP/TP of 0.1) while preserving a TPR of up to 0.88 (see brown line, Fig 4F). Hyperedge bundling is thus superior to strict thresholding in attenuating FP/TP with little decrease in TPR.

Importantly, our simulations show that the raw edges with largest correlational estimates might not correspond to the strongest or most important neurophysiological connections, because these estimates appeared to be correlated with reconstruction accuracy *(Supplementary).* The reconstruction accuracy is heterogeneous across source space, meaning high accuracy of sources may positively bias the *iPLV* estimates. This bias is another reason for including weak observations in FC graphs.

### Control parameters of hyperedge determine resolution and the balance among FP/TP, TPR, separability

In the current implementation, hyperedge bundling is controlled by the cutoff limit (CL) and the hyperedge size threshold (*θ_HEsize_*). CL determines the resolution of the hypergraph and the balance between noise (FP/TP) and *separability* of true hyperedges. Low CL values lead to low noise in hypergraphs but poor separation of true raw edges into distinct hyperedges. *θ_HEsize_* can be used to prune the smallest hyperedges to further reduce noise, albeit at a cost of pruning TP hyperedges.

We compared two clustering methods, UPGMA and Louvain. While the results showed clearly that by and large both clustering methods yielded similar performance, each method had interesting advantages. Louvain yielded better TPR than UPGMA for CL values between 0.05 and 0.25 (see Fig. 5B), and lower noise when singleton hyperedges were excluded (see Fig. 5A). UPGMA, on the other hand, yielded better separability of TP hyperedges throughout the control parameter ranges. Overall, using either clustering method with CL = 0.15-0.25 and *θ_HEsize_* = 1-2 will yield a large reduction in FP/TP (from 4 to 0.1-0.2) with good separability and negligible reduction in TPR.

In applications to real data where the truth graph is unknown, choosing parameters, *i.e.*, to control the trade-off between suppressing noise and maintaining high TPR and separability, can be based on both our simulation results and objectives of the research. If the objective of the hypothesis requires good separability (*e.g.*, establishing connectivity between specific visual areas to inferior parietal region), one should create high resolution hypergraphs, but this will be accompanied by sub-optimal noise reduction. Conversely, if the objective is to establish connectivity between the visual and parietal regions, a low resolution hypergraph (with low noise) is pertinent.

### Comparison of hyperedge bundling and symmetric orthogonalization

Symmetric orthogonalization is a pioneering solution to the overall problem of SIs in the context of amplitude correlation estimation (Colclough et al., 2015). Its predecessor, pairwise orthogonalization (Brookes et al., 2012; Hipp et al., 2012) excluded instantaneous mixing and evaluated amplitude correlations for each time-series pair at a time. It is thus applicable to the estimation all-to-all amplitude correlations similarly to any other bivariate AI-free metric for phase or other forms of coupling, and also suffers from SIs in the same manner (Palva et al., 2017).

Symmetric orthogonalization overcomes the problem of SIs by simultaneously removing zero phase-lag components from all source time series through a gradient descent procedure known as the Löwdin orthogonalization (Everson 1999; Löwdin 1950). Next, all-to-all amplitude correlations are estimated with partial correlation of amplitude envelopes to keep direct and remove indirect interactions (Marrelec et al., 2006). Because the partial correlation matrix is expected to be sparse, a graphical lasso regularisation of the inverse covariance matrix is applied to penalize near-zero elements (Banerjee et al., 2008; Friedman et al., 2008), which reduces noise in the partial correlation graph.

Symmetric orthogonalization effectively attenuates SIs caused both by signal leakage and by indirect true couplings (i.e., A ↔ C correlation, when true correlations are A ↔ B ↔ C). The two limitations of this method are: *i*) it is applicable only to the estimation of amplitude correlations, *ii*) it is limited by the rank of the data due to its dependence on singular value decomposition. For MEG/EEG data that are preprocessed with signal space separation (SSS) and temporal SSS methods, the rank of the data (~degrees of freedom) is often networks with less than 60-70 independent sources, such as the 19 regions per hemisphere used in (Colclough et al., 2015). For studying FC with greater parcellation resolutions (≫ 70) or with interaction metrics other than amplitude correlations, hyperedge bundling thus provides an alternative method for SI suppression. The similarities and differences between symmetric orthogonalization and hyperedge bundling are summarized in Table 1.

### Optimal source space for brain connectivity analyses

There are numerous MEG/EEG source reconstruction methods and the choice of method may have profound impacts on source connectivity analysis due to their difference in sensitivity to various synchronization profiles of the interacting sources (Hincapié et al., 2017). Although in the present study we used linear inverse operators (Hamalainen and Sarvas 1989; Hamalainen and Ilmoniemi 1994; Lin et al., 2006), hyperedge bundling can also be used with other source reconstruction methods as long as the amount of mixing among the sources/parcels can be quantified.

Parcel numbers in current MEG/EEG source connectivity studies range from tens of parcels, *e.g.*, 38 in (Colclough et al., 2015) and around 70 in (Farahibozorg et al.,; Hillebrand et al., 2012), up to 200-400 parcels(Lobier et al., 2017; Siebenhuhner et al., 2016; Zhigalov et al., 2017). We propose that the source-space for FC studies should have a fine spatial resolution that enables the separation of nearby independent signals to an extent allowed by the source reconstruction approach. Neither the neuronal source constellations nor the degrees of freedom in the data are likely to match any *a priori* chosen parcellation scheme and hence coarse parcellations can misrepresent or miss source areas that fall in between the parcels or are much smaller than the parcels.

Our approach to use 400 parcels aims to eliminate the possibility of such pitfall. Moreover, with finegrained parcellations, hyperedge bundling can well measure the mixing among raw connectivity edges and produce hypergraphs with high confidence of capturing and separating true interactions. Furthermore, the nodal groups connecting hyperedges can be utilized to coarsen a fine-grained source space in a data-driven manner and with consideration of the constraints posed by the source model. On the other hand, hyperedge bundling will likely to fail in a source-space of low spatial sampling, where the mixing similarity between observed edges is likely to be low due to initial low mixing among neighboring parcels.

## 6 Acknowledgements

We thank Prof. Karim Jerbi for comments to the work. We thank Dr. Giles Colclough for sharing source code and his insights to the symmetric orthogonalization method. This research was funded by the Academy of Finland grants 266745 and 281414. The funding bodies had no role in the design, data acquisition and analysis, decision to publish, or preparation of the manuscript.

